# Subways, zoning and social factors influence the dispersal and genetic connectivity of an iconic urban pest, the brown rat (*Rattus norvegicus*) across a complex cityscape

**DOI:** 10.1101/2025.04.26.650778

**Authors:** Matthew Combs, Jason Munshi-South

## Abstract

Cities present unique ecological challenges and opportunities for species that have adapted to human-dominated environments. To understand how urban landscape heterogeneity shapes the movements of an iconic urban animal, the brown rat (*Rattus norvegicus*), we investigated the factors influencing rat genetic connectivity across Manhattan, New York City. We generated genome-wide SNP data from rats sampled throughout Manhattan as well as paired surface-subway locations, created a habitat suitability model using systematic municipal rat survey data, and employed landscape genetic approaches to identify urban features influencing rat gene flow. Our models revealed that underground subway tunnels were the strongest facilitator of rat gene flow, with municipal zoning also strongly influencing connectivity, particularly through low / medium-density residential areas. Habitat suitability analysis showed that high human population density, old building ages, and lower median household income were the strongest predictors of rat presence. Subterranean sampling revealed that rats in subway stations exhibited greater genetic relatedness than surface rats at short distances but lower inbreeding coefficients, suggesting underground tunnels facilitate longer-distance dispersal while maintaining some connectivity to surface populations. These findings demonstrate that physical infrastructure, political, and socioeconomic factors shape urban rat ecology, with implications for more targeted approaches to pest management in complex urban environments. This study is the first comprehensive analysis of rat dispersal and genetic connectivity in relation to complex cityscape heterogeneity.

## INTRODUCTION

Over the last several decades, ecologists and evolutionary biologists have increasingly recognized that cities are valuable ecosystems that exhibit unique properties^1^. Urban ecologists have documented novel species assemblages^2^ and interactions^3^ in cities due to coexistence of the regional native species pool and myriad introduced taxa. Urban impacts on native biodiversity are heterogeneous. For example, plant^4^, bird^5^, and mammal^6^ diversity may be high in moderately urbanized areas, but invertebrate diversity tends to decrease substantially with urbanization^7^.

Cities are often located in areas with historically high levels of biodiversity, but are also often major centers of international trade and transport (particularly coastal shipping ports). These transport networks, challenging conditions, and novel infrastructure promote invasion of urban ecosystems by nonnative species. The most successful of these invasive species have evolved a substantial degree of dependence on humans^8^ to take advantage of our food supplies, waste streams, and urban infrastructure.

Three commensal rodents in particular, the house mouse (*Mus musculus*), black rat (*Rattus rattus*), and brown rat (*Rattus norvegicus*) are now globally distributed urban pests. *R. norvegicus* alone may have caused as much as $156 million in damage in the USA over the past several decades^9^. Problems associated with *R. norvegicus* infestations are accelerating, and urban municipalities are spending large sums of money and human capital on managing brown rat populations^10^. Long-term solutions to rat infestations have been elusive due to a poor understanding of commensal rat biology that can be applied in cities. Knowledge of how rats use urban resources and move through cityscapes has been particularly lacking, leading to reliance on reactive poisoning and trapping that is largely ineffective at the population level. Ecological investigations of real-world rat infestations are sorely needed to change this situation. **In this study, we use landscape genetic and habitat suitability models to examine the influence of urban infrastructure, natural habitat features, socioeconomic and political phenomena on rat presence, movements, and genetic structure in Manhattan, New York City, USA.**

Wildlife responds to urban heterogeneity in diverse ways, but researchers have tended to assume a traditional conservation paradigm where native species experience habitat fragmentation, ultimately leading to genetic differentiation and loss of genetic diversity^11^. However, it is clear that some species respond positively to urbanization, even if little is known about how these species interact with urban landscapes^12^. These successful species have been variously classified as urban exploiters^13^, synurbic^14^, or anthrodependent^8^ when their success in cities is predicated on access to human resources. Anthrodependent species are likely to interact with a wider array of environmental conditions in cities than native species that are facultative urban residents. Studies of anthrodependent species such as rats are also more likely to be generalizable across cities because these taxa are globally invasive, and thus may now or in the future be considered “native” to cities^15^. Urban rats also exert a significant negative toll on economies and public health because they damage infrastructure and food supplies and are reservoirs of zoonotic disease^16^. Understanding rat ecology and evolution is needed for urban risk management. Urban rats are particularly challenging to manage because they may use complex subterranean infrastructure such as subway and sewer systems in addition to typical terrestrial habitats. We leverage large spatial datasets describing the Manhattan landscape, habitat suitability modeling, and hundreds of rats genotyped at thousands of genome-wide single nucleotide polymorphisms (SNPs) to understand the connectivity of urban rats in a complex cityscape.

Resource availability, particularly human food subsidies, influences local abundance of anthrodependent species such as rats^17^. Landscape permeability to movement influences genetic connectivity, population persistence, and colonization dynamics in complex ways. Dozens of population genetic studies have documented the influence of urbanization on genetic drift and gene flow, but the magnitude and direction of these effects varies widely across species and cities^18^. For example, white-footed mice (*Peromyscus leucopus*)^19^ and redback salamanders (*Plethodon cinereus*)^20^ exhibit little connectivity across urban landscapes, but gene flow of black widow spiders (*Latrodectus mactans*)^21^, pigeons (*Columba livia*)^22^, and common wall lizards (*Podarcis muralis*)^23^ is facilitated by urbanization. Landscape genetic analyses take the extra step of modeling the influence of individual landscape variables on connectivity. These studies have largely focused on land cover classes, buildings, and transportation infrastructure to identify factors that restrict gene flow^12,24,25^. However, urban landscapes are composed of a complex mix of above- and below-ground infrastructure as well as remnant natural land cover and managed green spaces. Animal movements may also be influenced by social, cultural, and political variation among human communities, but these factors have typically not been incorporated into urban ecology and evolution studies^26,27^. Brown rats are likely affected by all of these factors, but it is currently unknown which landscape factors exert the greatest influence on rat population connectivity.

Logistical challenges and data limitations have hampered our ability to understand how rats use urban landscapes. Habitat suitability modeling, typically based on correlations between presence and ecological variables, requires high-confidence occurrence data^28^. City agencies rarely collect rat presence data using standardized methods, and instead rely on eyewitness reports from the public where species identity is unconfirmed and the probability of reporting is highly variable between neighborhoods. Landscape genetic analyses perform best when sampling of individuals can be conducted across entire landscapes^29^. Careful sampling design avoids bias from oversampling of relatives and facilitates analysis of the entire environmental gradient experienced by rats in a particular city. Targeted sampling around landscape features of interest may also be needed to provide insight into complex features, such as three-dimensional subterranean architecture that cannot be adequately captured using two-dimensional landscape rasters. A comprehensive landscape genetic analysis of an urban rat population has not previously been attempted to our knowledge. In this study, we sample rats across the entire island of Manhattan, NYC, build a habitat suitability model using thousands of presence data points from standardized surveys of rats in Manhattan, and incorporate sampling from subway stations as well as the terrestrial surface.

*Rattus norvegicus* are a powerful system for investigating the impacts of urbanization on landscape connectivity despite the challenges in studying them. Brown rats are highly dependent on human food resources, respond strongly to changes in human activity and the physical / social landscape ^30,31^, occur in most cities around the world, and are relatively easy to sample in large numbers. Short generation times and lifetime dispersal distances of brown rats can also produce spatial population genetic structure relatively quickly. Brown rats have the capacity to travel several km, but most stay within dozens of meters of where they were born^32–34^. Previous population genetic analyses in multiple cities indicate that both discrete barriers such as waterways, and diffuse quasi-barriers such as areas with poor habitat quality, are associated with population genetic structure at citywide scales^35^. Barriers such as major roads may also limit movement and create genetic structure at the scale of individual city blocks^33^. The most detailed field studies of rat movement also found that rats rarely move more than a few city blocks^31^.

New York City is in many ways an ideal place to study the influence of urban landscapes on rat connectivity and genetic structure. Brown rats were present in NYC no later than the American Revolutionary War^36,37^, and were acknowledged as a major problem in southern Manhattan by the early 19th century^10^. NYC is the most densely populated major city in North America and has been colloquially referred to as the “Ratopolis” due to its rat-friendly attributes. The 8.8 million permanent human residents and millions of daily commuters and tourists in NYC produce large amounts of food waste that ends up as garbage on the street or subway, or placed outside nightly in plastic bags for disposal. Rat-resistant containerization of garbage in New York City has remained an elusive goal. This constant stream of food, along with NYC’s subtropical humid climate^38^, promotes rat reproduction during all but the coldest months of the year. NYC also contains substantial harborage for rats in the form of subterranean infrastructure such as sewers and subway tunnels, as well as aging buildings and many formal and informal green spaces.

NYC agencies do not collect long-term data on rat population sizes, but civilian complaints indicate that rat infestations have been persistent and growing in recent years^39^. Statistical modeling of rat presence data from a one-time NYC public health survey identified residential buildings, older buildings, parks, schools, and rail or subway lines as correlated with rat presence, and improved waste management as associated with decreased rat presence^40^. A similar analysis of public health inspections triggered by civilian complaints found subway lines, parks & recreational lands, older residential buildings and vacant housing as predictive of rat sightings^41^. These valuable results have identified correlates of high-quality rat habitat in NYC, but do not explain what factors may promote or hinder the dispersal of rats through heterogeneous urban landscapes. In previous population genetic studies in Manhattan^32,35^, we found that pairs of rats exhibit high relatedness at distances less than 200 meters, with relatedness greater than zero extending out to 1.4 km. Gene flow was not evenly distributed across the landscape, however. An area of low migration rates and high inbreeding coefficients in midtown Manhattan has created two evolutionary clusters in northern (uptown) and southern (downtown) portions of the island. This earlier study did not identify specific landscape attributes that influence gene flow, however. Many outstanding questions about urban rat presence and movement still remain, such as whether rats in subterranean habitats comprise local population isolates or whether rats move freely between the surface and underground.

Here, we 1) generate and compare landscape genetic models that examine rat connectivity across the Manhattan cityscape in relation to natural land cover, the built environment, and human social variation; 2) construct a habitat suitability model for rats and examine whether it predicts rat connectivity; and 3) examine genetic relatedness of rats collected in subway tunnels and nearby surface environments. This study is the first comprehensive analysis of rat dispersal and genetic connectivity in relation to urban landscape heterogeneity.

## METHODS

### Study Site and Sampling

A full accounting of our main sampling scheme and study site has been presented previously in Combs et al. (2018)^32^. New data collection and analyses are detailed below. In brief, we used lethal snap traps to capture 393 brown rats across the island of Manhattan, New York City, NY between June 2014 and December 2015 (**Figure 1a**). We sampled tail tissue from deceased rats and stored them in 70% ethanol for downstream DNA analyses. Trapping was conducted on municipal land and private property with permission of landowners, and was targeted towards areas with recent signs of rat burrowing or other activity (e.g. fecal droppings, sebum trails, footprints, or sightings of dead or live rats). We limited our sampling to Manhattan because we could feasibly sample the entire island, there is excellent information available on the attributes of the Manhattan landscape, and Manhattan has had a thriving rat population for decades due to an abundance of food, water and harborage available to rats^10^. All animal handling procedures were approved by the Fordham University IACUC.

**Fig. 1:**
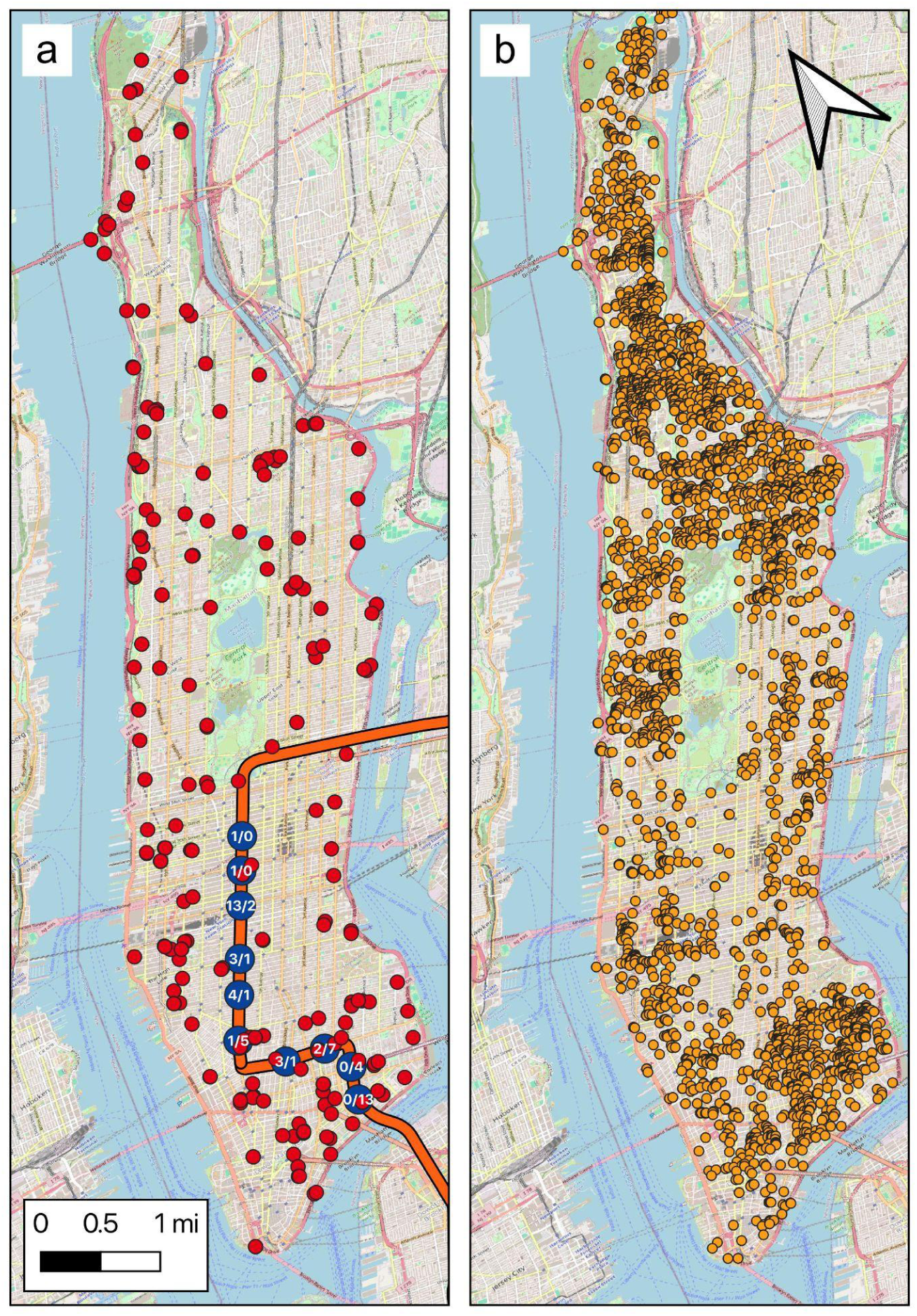
Individual rats sampled for genetic analyses across Manhattan and in subway stations, and independent inspection points that detected active rat signs. a Red dots represent individual rats trapped for genetic analyses at the surface level in Manhattan. The orange line represents a subway line where rats were trapped within and around subway stations (blue dots). Numbers within blue dots represent above / below ground sample sizes for each station. b Orange dots represent active rat signs detected by the NYC Department of Health in a systematic survey of every property in Manhattan in 2012. These locations were used as presence points for habitat suitability modeling.

### Generation and processing of genomic data

We produced genomic libraries from Manhattan rats using the ddRADSeq protocol of Peterson et al. (2012)^42^. Full details of procedures for generating population genomic data appear in Appendix S1 of Combs et al. (2018)^32^. We sequenced these individually barcoded libraries using paired-end 125 bp sequencing on three Illumina HiSeq 2500 lanes. All of these sequencing reads are available on the NCBI SRA under accession PRJNA414893. We then demultiplexed sequence reads by individual using STACKS v1.35^43^, and aligned each individuals’ reads to the Rnor6.0 version of the *Rattus norvegicus* reference genome^44^ using BOWTIE2 with default settings^45^. We then used STACKS to call and filter SNP genotypes. We filtered out loci sequenced in fewer than 75% of individuals, applied a minor allele frequency cutoff of 2.5%, and removed any individual with greater than 50% missing SNP genotypes. After applying these filters, we retained 262 individuals genotyped at 61,401 polymorphic loci. We then used the program BED2DIFFS v1 in the EEMS software package^46^ to generate a pairwise matrix of average genetic dissimilarity across nonmissing loci for downstream landscape genetic analyses.

### Landscape data and resistance surfaces

We generated GIS raster layers of landscape data to test specific hypotheses for how features of the urban landscape influence gene flow of urban brown rats (**Table 1**). These hypotheses relate to three major categories of landscape variation: 1) the non-built physical landscape; 2) the built physical landscape; and 3) the human social landscape. The non-built physical landscape describes natural landcover classes and geologic structure of the environment that influence resource availability and potential impediments or facilitators of movement. The built physical landscape describes anthropogenic landcover classes, infrastructure, and building characteristics that rats may exploit for resources and dispersal opportunities. The social environment describes non-physical aspects of the environment that encompass the behavior and organization of human activity, which may in turn influence the habitat quality or movement potential in a landscape.

**Table 1.**
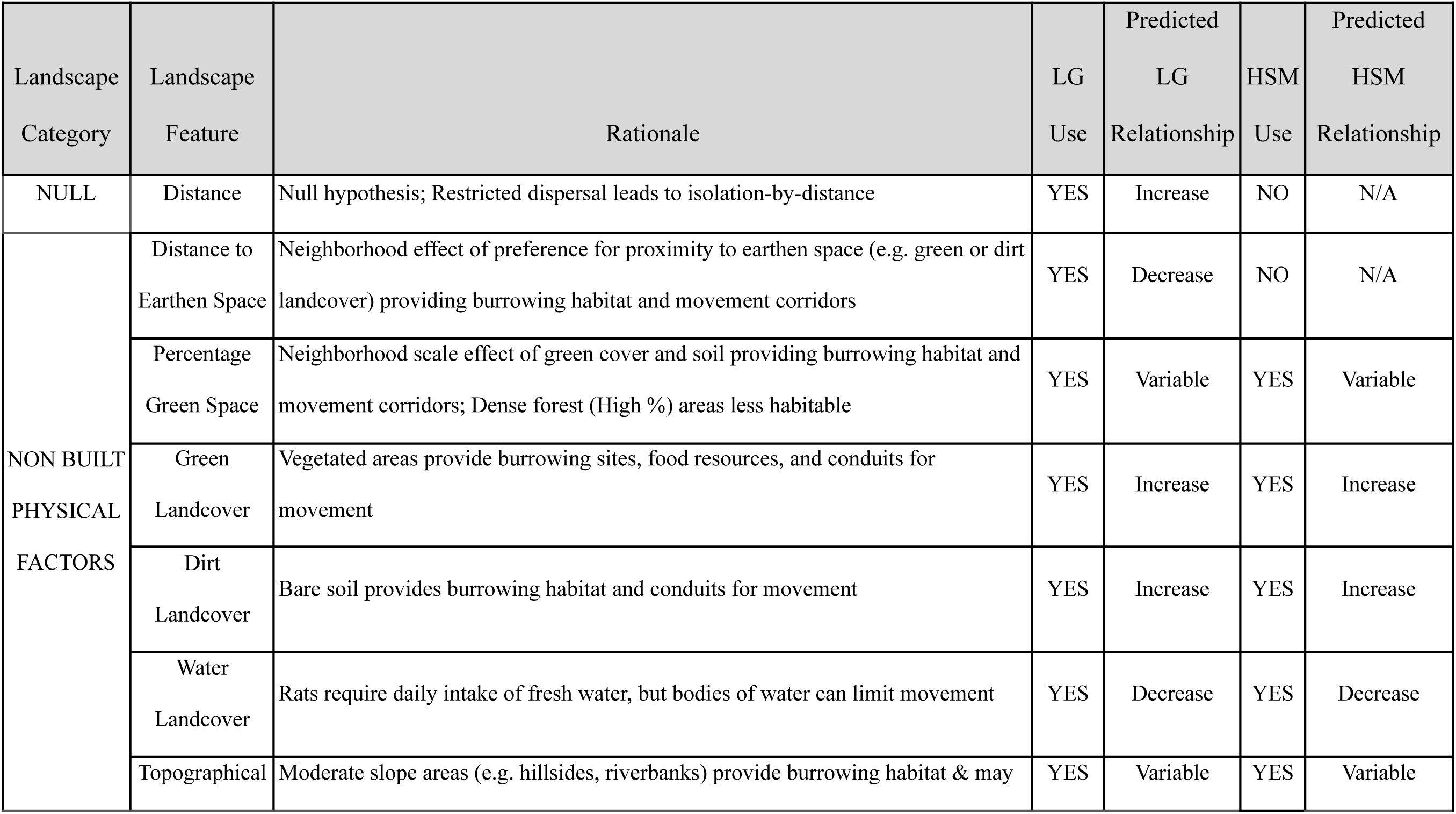
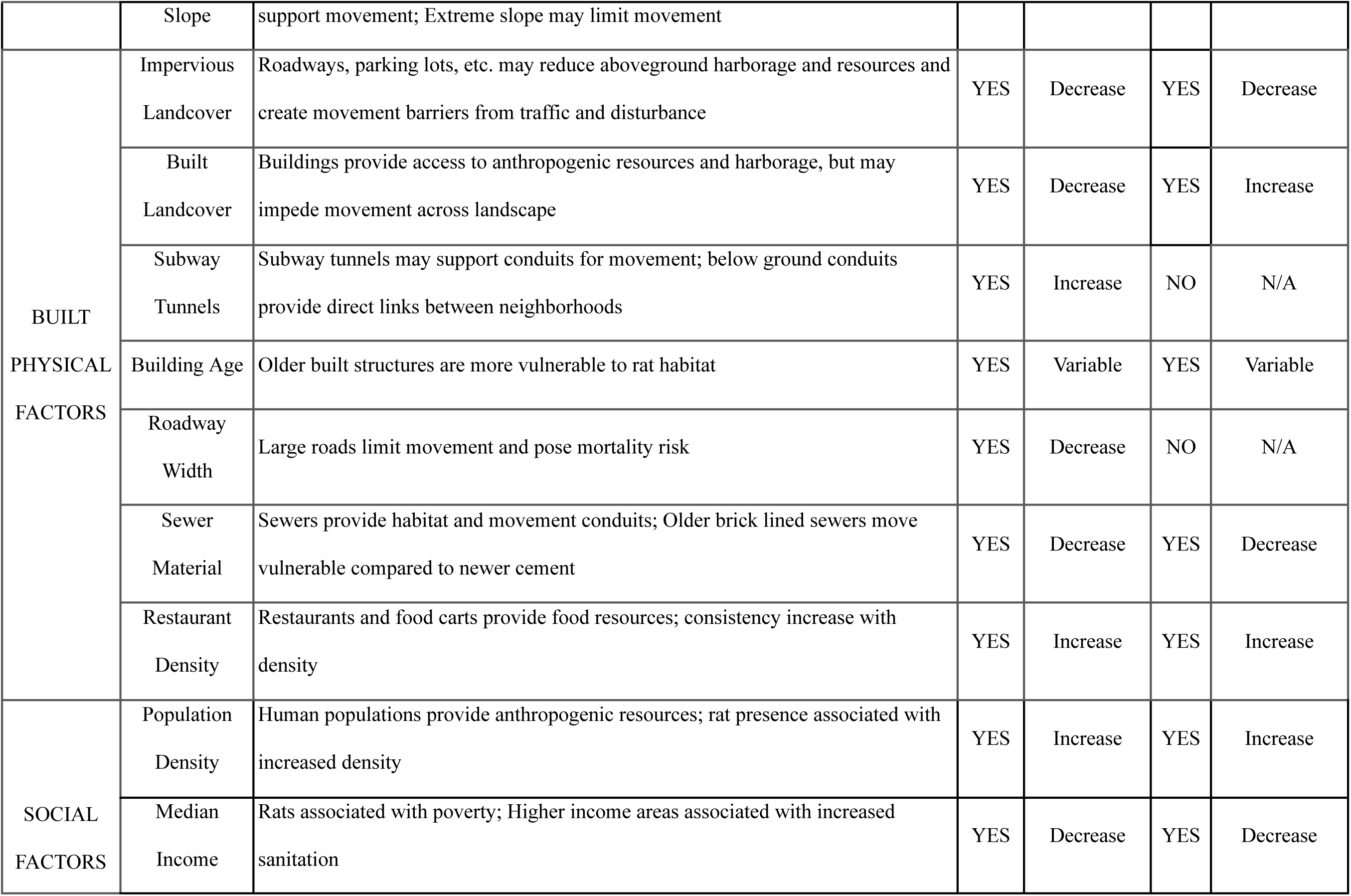
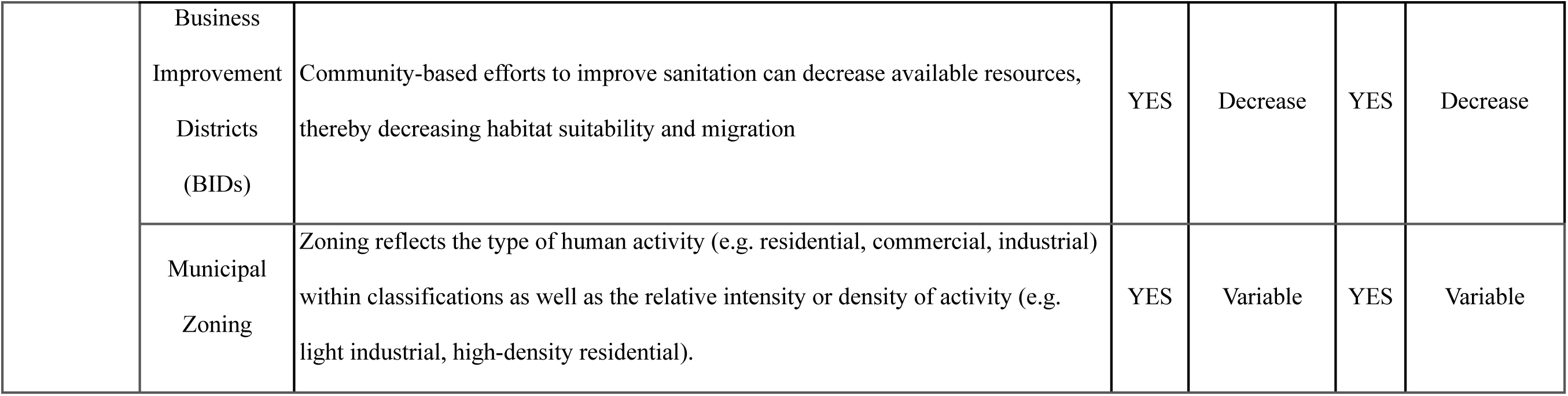
Landscape variables, rationale for inclusion in landscape genetic models, and inclusion / predictions for direction of effects on genetic connectivity (LG) and habitat suitability (HSM).

| Landscape Category | Landscape Feature | Rationale | LG Use | Predicted LG Relationship | HSM Use | Predicted HSM Relationship |
| --- | --- | --- | --- | --- | --- | --- |
| NULL | Distance | Null hypothesis; Restricted dispersal leads to isolation-by-distance | YES | Increase | NO | N/A |
| NON BUILT PHYSICAL FACTORS | Distance to Earthen Space | Neighborhood effect of preference for proximity to earthen space (e.g. green or dirt landcover) providing burrowing habitat and movement corridors | YES | Decrease | NO | N/A |
|  | Percentage Green Space | Neighborhood scale effect of green cover and soil providing burrowing habitat and movement corridors; Dense forest (High %) areas less habitable | YES | Variable | YES | Variable |
|  | Green Landcover | Vegetated areas provide burrowing sites, food resources, and conduits for movement | YES | Increase | YES | Increase |
|  | Dirt Landcover | Bare soil provides burrowing habitat and conduits for movement | YES | Increase | YES | Increase |
|  | Water Landcover | Rats require daily intake of fresh water, but bodies of water can limit movement | YES | Decrease | YES | Decrease |
|  | Topographical | Moderate slope areas (e.g. hillsides, riverbanks) provide burrowing habitat & may | YES | Variable | YES | Variable |
|  | Slope | support movement; Extreme slope may limit movement |  |  |  |  |
| BUILT<br>PHYSICAL<br>FACTORS | Impervious<br>Landcover | Roadways, parking lots, etc. may reduce aboveground harborage and resources and create movement barriers from traffic and disturbance | YES | Decrease | YES | Decrease |
|  | Built<br>Landcover | Buildings provide access to anthropogenic resources and harborage, but may impede movement across landscape | YES | Decrease | YES | Increase |
|  | Subway<br>Tunnels | Subway tunnels may support conduits for movement; below ground conduits provide direct links between neighborhoods | YES | Increase | NO | N/A |
|  | Building Age | Older built structures are more vulnerable to rat habitat | YES | Variable | YES | Variable |
|  | Roadway<br>Width | Large roads limit movement and pose mortality risk | YES | Decrease | NO | N/A |
|  | Sewer<br>Material | Sewers provide habitat and movement conduits; Older brick lined sewers move vulnerable compared to newer cement | YES | Decrease | YES | Decrease |
|  | Restaurant<br>Density | Restaurants and food carts provide food resources; consistency increase with density | YES | Increase | YES | Increase |
| SOCIAL<br>FACTORS | Population<br>Density | Human populations provide anthropogenic resources; rat presence associated with increased density | YES | Increase | YES | Increase |
|  | Median<br>Income | Rats associated with poverty; Higher income areas associated with increased sanitation | YES | Decrease | YES | Decrease |
|  | Business Improvement Districts (BIDs) | Community-based efforts to improve sanitation can decrease available resources, thereby decreasing habitat suitability and migration | YES | Decrease | YES | Decrease |
|  | Municipal Zoning | Zoning reflects the type of human activity (e.g. residential, commercial, industrial) within classifications as well as the relative intensity or density of activity (e.g. light industrial, high-density residential). | YES | Variable | YES | Variable |

The resulting rasters were used as inputs for habitat suitability modeling and / or converted to resistance values for landscape genetic modeling (see sections below for details). Though some data were available at finer resolution, we resampled all layers to 30 m^2^ resolution for computational feasibility. This resolution is in line with the scale at which rats typically experience the landscape^34,47^. Details on the data sources and construction of each landscape raster are included in the supplementary materials (Table S1). Prior to downstream analyses, we tested for multicollinearity among landscape rasters using the Band Collection Statistics tool in ARCMAP v10.6, which calculates Pearson’s correlation coefficient between all pairs of rasters. No layers were found to be highly correlated, and thus all layers were retained for analyses.

### Habitat Suitability Modeling

To perform habitat suitability modeling, we first identified a set of high-confidence *R. norvegicus* locations from a 2012 survey of active rat signs (ARS) conducted by the New York City Department of Health and Mental Hygiene across every property parcel in Manhattan (**Figure 1b**). We extracted the centroid of each parcel with confirmed ARS and combined these points with locations of rats that we trapped for this study (**Figure 1a**). We then used a suite of landscape variables (**Suppl. Methods, Tables S1-S2**) to examine the influence of urban heterogeneity on rat presence. To this end, we used MAXENT v3.4^48^ to develop a habitat suitability map. Maxent predicts species occurrence likelihood by linking presence-only location data with rasters of landscape attributes, and has been shown to perform well across a wide variety of conditions^49^. To create the model, we first optimized Maxent’s regularization parameter, which penalizes for model complexity, using the model selection procedure available through ENMTools^50^. We tested 20 regularization coefficients spaced evenly between values from 0.1 – 2.0 across independent model runs and then selected the best-fitting model based on AICc scores. Using the optimized parameter, we performed 10 Maxent iterations, specifying auto features and using cloglog output. We iteratively removed the layer with the lowest contribution until AICc increased, which led to the removal of brick sewer density and topographical slope from the model.

### Landscape genetic modeling

We used individual-based isolation-by-resistance (IBR) modeling to quantify the role of urban landscape heterogeneity on *R. norvegicus* gene flow using the ResistanceGA^51^ optimization framework. This method parameterizes each resistance surface through a genetic algorithm procedure, removing the need for assigning resistance values based on expert opinion. The approach explores parameter space using a series of transformations (varying both shape and maximum value parameters) applied to the original raster values, which aim to improve model fit between pairwise genetic distances and resistance distances. Resistance distances were calculated through the commuteDistance function in the gdistance^52^ package in R, which is nearly identical to resistance distance calculations in Circuitscape^53^, but with improved computation time. We used log likelihood as the objective function during optimization, which implemented linear mixed-effect models (LMMs) where genetic distance was the dependent variable and landscape resistance was the fixed effect. We used maximum likelihood population effects (MLPE) parameterization to define random effects that account for non-independence between genetic and resistance distances^54^, as implemented in the lme4^55^ package in R.

Using the citywide set of all genotyped samples, we first independently optimized each landscape variable hypothesized to influence rat gene flow independently using ResistanceGA’s *ss_optim* function. We also established multivariable sets of landscape layers hypothesized to influence functional connectivity in combination, which we optimized using ResistanceGA’s *ms_optim* function. These multi-layer sets included all built features, all non-built features, all social features, naturally available resources of green and water landcover, anthropogenic food resources including restaurant and population density (i.e. Rat Food), all transportation infrastructure, subway and sewer infrastructure, subway infrastructure and municipal zoning classes, income and green landcover or percentage of greenspace together, population density and green landcover or percentage of greenspace, and income and population density together . See **Table 1** for a description of each variable and their groupings. Following optimization, we checked for normality of residuals and overall linear relationship between genetic distance and optimized resistance distance to ensure model fit. We used AICc to rank the fit of all models.

### Population genomics of subway rats

To better understand the role of underground transportation tunnels on the genetic connectivity of Manhattan rats, we collected additional rat tissue samples along a portion of a single subway line (the F train) in Manhattan from July to November 2016. We used a paired sampling approach, trapping rats both below-ground in each subway station between the 47th Street and East Broadway stations as well as above-ground within one city block of each sampling station (**Figure 1a**). These ten stations are arrayed along a roughly linear track that is ∼5.5 km long. Rat trapping, DNA extraction, ddRAD library preparation and sequencing, and SNP calling procedures for these samples were identical to those described above for citywide sampling. We generated a dataset of 32,513 SNPs for the above- and below-ground comparison. After removing samples with missing data greater than 50%, we retained 25 below-ground rats and 32 above-ground rats for downstream analyses. We examined basic population genetic summary statistics for above- and below-ground rats separately. We also used the resulting SNP dataset to generate matrices of genetic dissimilarity averaged across non-missing loci using the *bed2diffs_v1* script from the EEMS software (Petkova 2016) for rats sampled above- and below-ground. We then plotted the relationship between genetic dissimilarity and geographic distance for above- and below-ground subway rats separately. Additionally we performed principal coordinate analysis using the combined set of all above- and below-ground rats.

## RESULTS

### Habitat Suitability Modeling

We developed a habitat suitability model for Manhattan to understand the ecological influence of urban landscape features on rat presence, using rat presence records from both a systematic municipal survey of property parcels and locations of rats we trapped across Manhattan (**Figure 1**). Predicted rat presence was highest in Manhattan’s southeastern neighborhoods (East Village, Lower East Side), and much of the Harlem neighborhoods stretching north / northeast. Smaller pockets of high-quality rat habitat were also predicted in the southwest (West Village) and central west (Hell’s Kitchen), while low habitat suitability was identified across much of central midtown, between the southeast and southwest neighborhoods, and in areas with forested green spaces (**Figure 2**).

**Fig. 2:**
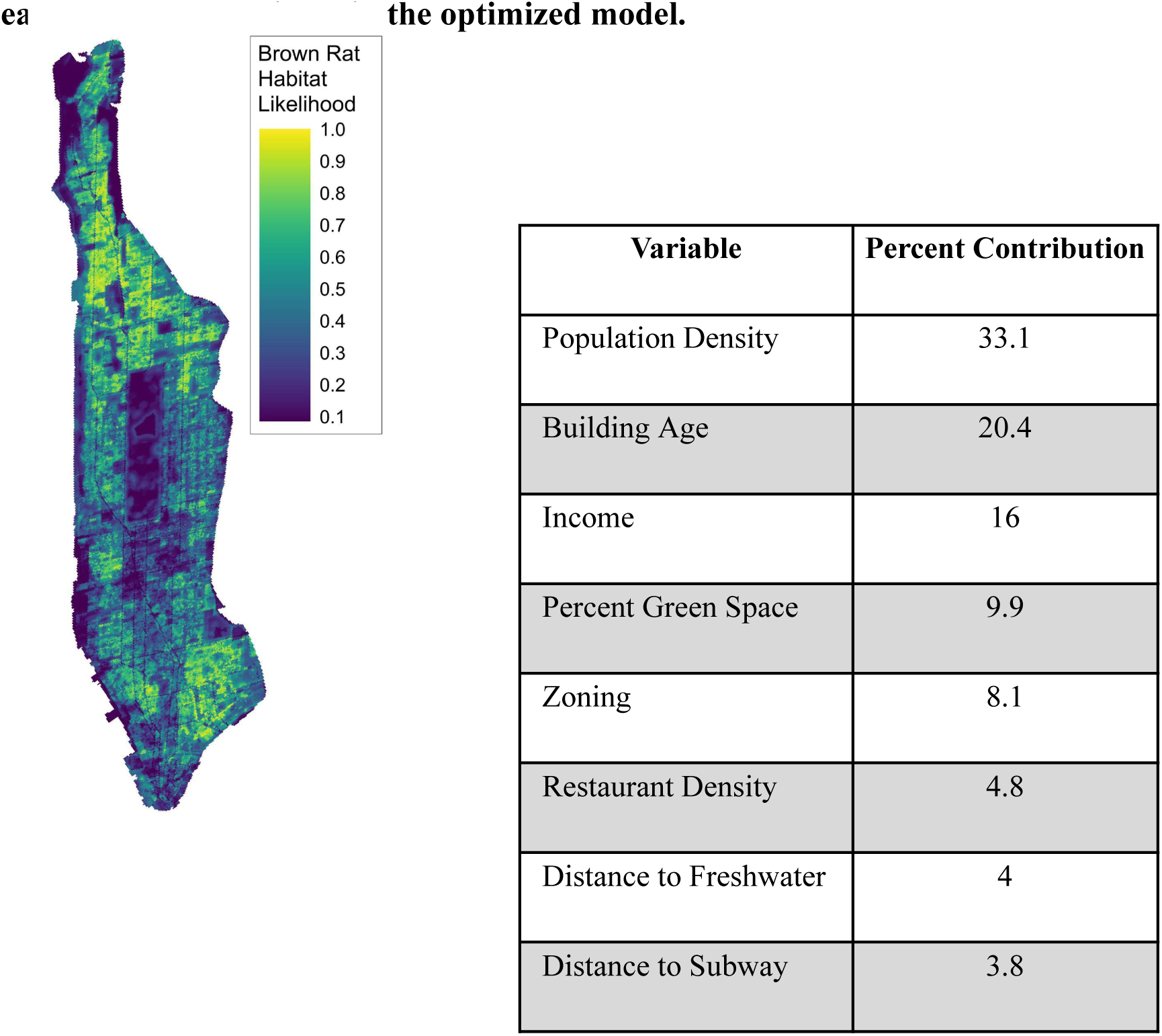
Habitat suitability model for brown rats in Manhattan and relative contribution of each landscape variable in the optimized model.

Population density, building age, and median household income were the strongest predictors of rat presence, contributing 33.1%, 20.4%, and 16.0% respectively to the final model (**Figure 2**). Habitat suitability was generally linked to more densely populated areas, with peaks at moderate densities, as well as buildings built around 1900 and census tracts with relatively low median income between $20,000 - $60,000 US per year (see **Figs. S1-S8** for response curves for each final variable). Conversely, regions of particularly low habitat suitability were linked to extremely low human population density, buildings built around 1950, and census tracts with relatively high median income above $75,000 US per year. The density of green space and neighborhood zoning had moderate influence on the model, contributing 9.9% and 8.1% respectively. Predicted rat presence was highest in areas with moderate green space coverage (10-30% of landcover) and in neighborhoods zoned for low / medium density residential development, followed by high density residential development. Habitat suitability decreased linearly with an increasing percentage of green landcover over 30% and was lowest in areas zoned for medium / heavy manufacturing. Distance to water and the nearest subway line were also included in the final model after variable reduction, but contributed only 4% and 3.8% respectively.

### Landscape Predictors of Citywide Genetic Connectivity

The presence of underground subway tunnels was the best predictor of rat genetic connectivity across Manhattan based on AICc scores of optimized IBR models (**Table 2**). We ran a second iteration with slightly different AICc scores and very similar rankings (Table S2). The final model indicated that subway tunnels are 635X less resistant to connectivity than non-subway areas, indicating that underground tunnel infrastructure facilitates rat gene flow across Manhattan. Many subway tunnels run along the island’s north-south axis, with more diffuse networks in the island’s northern uptown neighborhoods. The subway network is more spatially dense approaching the island’s southern downtown neighborhoods (**Figure 3A,B**). Municipal zoning classifications also showed strong predictive value for citywide gene flow, ranking second among univariate models and present in three of the five top performing models. Low to medium-density housing and commercial areas facilitated rat gene flow. These areas of high rat connectivity include Manhattan’s central and eastern Harlem neighborhoods and a swathe connecting uptown and downtown neighborhoods along the eastern side of the island.

**Fig. 3.**
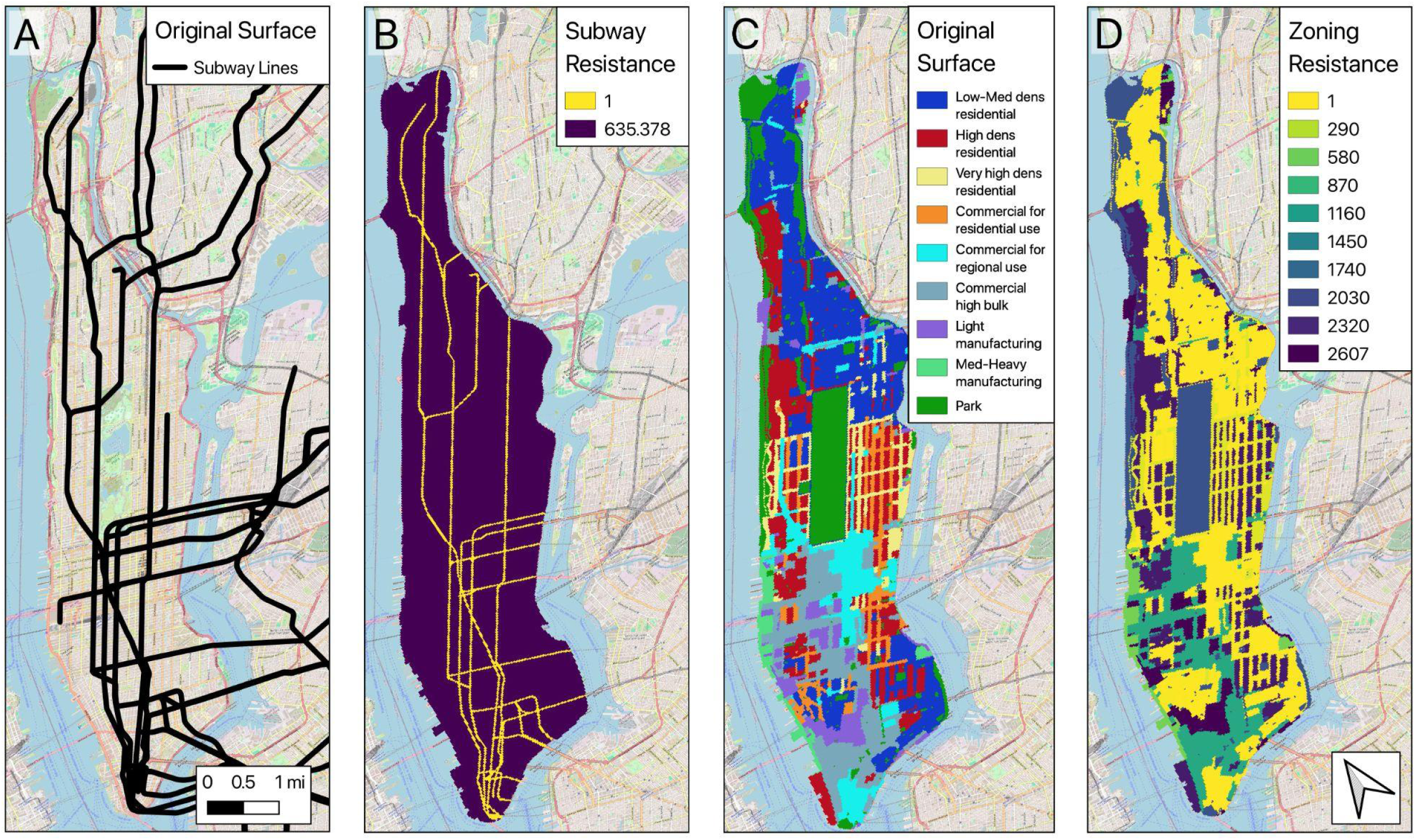
Original landscape datasets and highly-ranked landscape resistance models for subway lines (A,B) and zoning (C,D) in Manhattan.

**Table 2.** Univariate and multivariate landscape genetic models ranked by corrected Akaike’s Information Criterion (AICc).

| Model | Type | Mean AICc<br>Rank | Std. Dev.<br>AICc Rank | AICc | Marginal R<sup>2</sup> | Conditional R<sup>2</sup> |
| --- | --- | --- | --- | --- | --- | --- |
| Subway Tunnels | Uni | 1 | 0.0 | -159814.9 | 0.594 | 0.964 |
| Subway & Sewer | Multi | 2 | 0.0 | -159808.6 | 0.594 | 0.964 |
| Municipal Zoning | Uni | 3.5 | 0.7 | -156480.3 | 0.646 | 0.971 |
| Subway & Zoning | Multi | 3.5 | 0.7 | -157698.6 | 0.646 | 0.974 |
| All Social | Multi | 5.5 | 0.7 | -152572.2 | 0.569 | 0.926 |
| Rat Food | Multi | 5.5 | 0.7 | -153027.4 | 0.623 | 0.977 |
| All Natural | Multi | 8 | 0.0 | -151823.5 | 0.608 | 0.970 |
| GreenLC & PopDen | Multi | 8.5 | 2.1 | -150320.7 | 0.606 | 0.968 |
| Landcover | Multi | 8.5 | 2.1 | -152048.7 | 0.586 | 0.952 |
| Roadway Width | Uni | 9 | 0.0 | -150426.2 | 0.583 | 0.955 |
| Distance to Earthen Space | Uni | 11 | 0.0 | -150173.2 | 0.511 | 0.920 |
| Natural Resources | Multi | 12 | 0.0 | -150086.7 | 0.616 | 0.971 |
| Green LC | Uni | 13 | 0.0 | -150086.3 | 0.616 | 0.971 |
| GreenLC & Income | Multi | 14 | 0.0 | -150080.1 | 0.616 | 0.971 |
| Percentage Green Space | Uni | 15 | 0.0 | -149711.2 | 0.455 | 0.895 |
| PercentGreen & PopDens | Multi | 16.5 | 0.7 | -149506.2 | 0.444 | 0.889 |
| All Built | Multi | 17 | 1.4 | -149000.9 | 0.250 | 0.767 |
| PercentGreen & Income | Multi | 17.5 | 0.7 | -149493.0 | 0.440 | 0.886 |
| Sewer Material | Uni | 19 | 0.0 | -148565.8 | 0.520 | 0.895 |
| Built LC | Uni | 20 | 0.0 | -148264.0 | 0.526 | 0.910 |
| Topographic Slope | Uni | 21 | 0.0 | -147761.7 | 0.266 | 0.780 |
| Impervious LC | Uni | 22 | 0.0 | -147748.8 | 0.521 | 0.930 |
| Building Age | Uni | 23 | 0.0 | -147580.8 | 0.612 | 0.969 |
| Habitat Suitability Model | Uni | 25 | 0.0 | -147218.5 | 0.536 | 0.923 |
| Income & PopDens | Multi | 25 | 1.4 | -147386.8 | 0.198 | 0.724 |
| Restaurant Density | Uni | 26.5 | 0.7 | -146709.4 | 0.490 | 0.864 |
| BIDs | Uni | 27.5 | 0.7 | -146515.8 | 0.544 | 0.909 |
| Population Density | Uni | 28 | 5.7 | -145571.4 | 0.161 | 0.706 |
| Water LC | Uni | 28.5 | 0.7 | -145681.7 | 0.165 | 0.709 |
| Median Income | Uni | 29.5 | 0.7 | -145648.4 | 0.173 | 0.714 |
| Dirt LC | Uni | 30.5 | 0.7 | -145638.5 | 0.200 | 0.738 |
| Distance | Uni | 31.5 | 0.7 | -145575.1 | 0.161 | 0.706 |

Conversely, high density residential areas, high-bulk commercial areas, industrial areas, and city parks were all predicted to strongly impede gene flow, resulting in several apparent barriers to rat dispersal particularly along the island’s western side both uptown and downtown, including between the East Village and West Village neighborhoods (**Figure 3C,D**).

Rankings for model performance showed little variation across the two independent optimization iterations, with most models showing a standard deviation of less than 1.0. The habitat suitability model showed relatively poor performance, ranking in the bottom third of all tested models. The distance-only model ranked last among all tested models, suggesting that each landscape variable explained rat connectivity more than geographic distance by itself.

### Genetic Connectivity of Subway and Surface Rats

Population genetic summary statistics among the two cohorts were similar (**Table 3**) with the exception of decreased inbreeding coefficients among underground samples and the number of private alleles. Aboveground samples exhibited 594 private alleles, while the underground samples exhibited only 1 private allele, strongly suggesting that rats found underground are sourced from the aboveground population and contain very few unique genetic variants.

**Table 3:** Population genetic summary statistics for aboveground and belowground rats sampled along the F-line of the NYC subway system. Standard deviations provided in parentheses.

| | $H_O$ | $H_E$ | $P_i$ | $F_{IS}$ | Private Alleles |
| --- | --- | --- | --- | --- | --- |
| Underground | 0.224 (0.024) | 0.247 (0.001) | 0.252 (0.001) | 0.105 (0.012) | 1 |
| Aboveground | 0.22 (0.016) | 0.258 (0.001) | 0.26 (0.001) | 0.144 (0.015) | 594 |

Plots of pairwise genetic distance versus geographic distance for the belowground and aboveground subsets indicate that rats sampled within the underground subway stations were more closely related on average than those collected at the city surface, particularly at short distances (**Figure 4**). Linear models fit to belowground and aboveground rats separately indicate that beyond ∼4km belowground rats are no more closely related than aboveground rats. This finding suggests that mating between rats from subway stations and surface habitats increases the potential for gene flow among the Manhattan rat population. Despite the increased relatedness of subway rats at short distances, they exhibit substantially lower inbreeding coefficients than aboveground rats, indicating that subway rats experience a relative increase in reproduction with genetically dissimilar individuals.

**Fig. 4:**
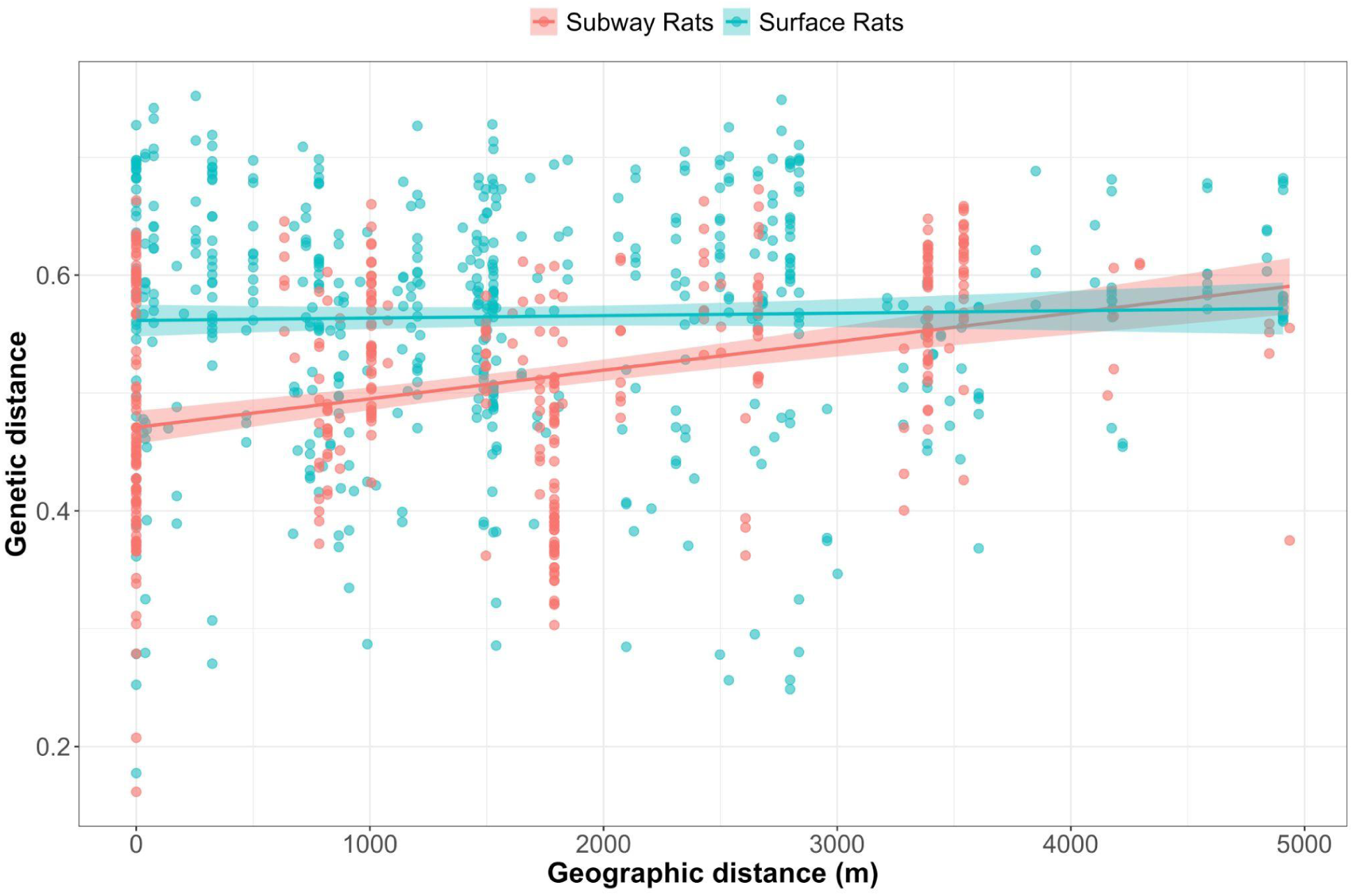
Pairwise geographic distance vs. pairwise genetic distance for brown rats sampled in subway stations (orange) vs. at the surface (blue).

## Discussion

In this study we show that the built environment, political factors, and socioeconomic variation in New York City influence the distribution and genetic connectivity of an archetypal urban species, the brown rat (*Rattus norvegicus*). While many studies have documented an influence of urbanization on gene flow or genetic drift^18^, very few have identified specific factors associated with the spatial distribution of genetic variation in cities. White-footed mice in NYC exhibit genetic connectivity between populations when tree canopy cover is sufficiently extensive^56^, and wall lizards in Germany exhibit greater gene flow when railway lines are present^23^. Movement of species that can’t fly is likely to be affected by the distribution of green or gray land cover in cities^57^, although the specific types of infrastructure that are important may vary widely.

White-footed mice are forest rodents that are behaviorally and morphologically adapted to moving through dense vegetation, whereas wall lizards favor dry, sunny, rocky habitats associated with railroads and stone walls. Brown rats typically build and occupy concealed earthen burrows, but use a wide variety of subterranean infrastructure like subways and sewer systems^58,59^ in cities around the world.

Urban ecologists have repeatedly called for incorporating social, economic and political variation into our analyses^26,27^, and some trends are now emerging. We found that zoning was one of the strongest predictors of genetic connectivity among urban rats. Zoning is a legal framework dictating what types of buildings can be built and for what human uses (e.g. residential, commercial, or industrial) in a particular area, and is often highly politicized. Similarly, several studies have identified historical redlining as a legacy factor that negatively influences biodiversity^60,61^, collection of community science data^62,63^, and even genetic diversity^64^. Redlining was a program in the USA to geographically delineate urban neighborhoods that should or should not receive real estate loans, resulting in disinvestment primarily in lower-income neighborhoods with an industrial presence. Political and social factors that concentrate poverty in certain urban neighborhoods are likely to have broad-ranging implications for other species, which will be detectable as a correlation with socioeconomic variables like median household income. For example, relative wealth of neighborhoods influences the strength of selection on goldenrod gall size in the Toronto metropolitan area, primarily detectable as greater bird predation in wealthier neighborhoods leading to directional selection for smaller galls^65^. Levels of biodiversity of many taxonomic groups are positively correlated with socioeconomic status, a phenomenon known as the “luxury effect”^66^. While income was not important in our landscape genetic models, it was one of the strongest variables predicting rat habitat quality, along with human population density and age of buildings. Not surprisingly, neighborhoods with many low-income residents inhabiting old buildings are most likely to have a high density of rats in NYC. While overall biodiversity may decrease with declining socioeconomic status, the presence and density of pests harmful to human health are likely to increase. In addition to rats, infestations of mosquitoes^67^, bed bugs^68^, cockroaches and house mice^69^ all exhibit such correlations. Effects may even be synergistic, as invasive mosquitoes have been found to feed heavily on rats in lower-income urban neighborhoods^70^.

Our study provides unique insight into how construction and management of complex urban ecosystems drives evolutionary processes for one of the most widely distributed and economically damaging pests worldwide. It remains an open question whether similar analyses in other cities would identify the same factors influencing rat genetic connectivity. However, population genomic comparisons of rats in Manhattan, New Orleans, Vancouver, and Salvador, Brazil indicate that patterns may differ across cities^35^. In that analysis, waterways (New Orleans) and roads (Vancouver and Salvador) were important in creating genetic structure except in Manhattan, where there was a genetic discontinuity created by a primarily commercial neighborhood^32^. Waterways were also important in creating genetic structure among rats in Baltimore^71^. Our finding that municipal zoning impacts genetic structure of rats helps explain the population genetic patterns we reported in our earlier population genetic analysis, particularly a split between uptown and downtown Manhattan associated with a commercial area in midtown Manhattan, as well as more subtle subpopulation differentiation between rats in Manhattan’s southeastern and southwestern primarily residential neighborhoods separated by high bulk commercial and light manufacturing areas^32^. To our knowledge, this study is the first to report impacts of zoning on population genetic structure of any species.

Zoning is a process through which a government entity divides land into “zones” that are defined by the uses, size of subdivided lots, and types and density of buildings that are allowed in the zone (among other possible restrictions). Land use regulations have a long history, but zoning became prevalent in cities in the United States in the early twentieth century^72^. The first comprehensive zoning law in the United States took effect in NYC in 1916^73^, but many other cities quickly followed suit. The 1916 NYC ordinance restricted the height (no more than 2.5X the street width), use, and lot coverage (i.e. footprint of buildings compared to lot size) of buildings, with the goals of avoiding shading and overcrowding caused by tall buildings^74^. The city was experiencing a real estate boom at the time, particularly along subway lines with the opening of the first stations in 1904. Previously, there were virtually no restrictions on the height and purpose of new buildings, and lucrative retail businesses and residential neighborhoods feared overcrowding and degraded quality of life from large manufacturing facilities. The contemporary landscape of Manhattan still reflects zoning decisions made decades ago, and our models captured the influence of these decisions on the genetic connectivity of the contemporary rat population. Not surprisingly, low- to medium-density residential and commercial areas facilitate genetic connectivity of rat populations in Manhattan. Neighborhoods zoned for these uses are likely to provide steady access to food resources from apartment buildings, restaurants, food carts and street litter from pedestrians. Subway stations also tend to be clustered in these neighborhoods, although they occur in other parts of the city as well. The NYC government inspection data that we used to build our HSM indicates that these areas have the most active rat signs, and thus are likely sources of rat dispersers that maintain genetic connectivity throughout residential areas. General agreement between independent inspection data and our landscape genetic modeling provides a higher degree of confidence in our results than the landscape genetic results alone.

In contrast to residential areas, high bulk commercial and manufacturing areas were identified by our modeling as factors that restrict rat connectivity in Manhattan. “High bulk commercial” refers to very large commercial properties with high ceilings, such as warehouses and distribution centers. Manufacturing facilities also tend to be located in similar types of large, open buildings, resulting in fewer interstitial spaces within and between buildings that rats can exploit for nesting and movement. These facilities may require relatively few workers, and if they are not used to store food then may not provide adequate resources to sustain large rat populations. Food warehouse / distribution centers are also likely to have enhanced pest management to protect consumable goods, and thus even these facilities may be relatively poor at sustaining rat populations. We also found that the highest-density residential areas were associated with reduced connectivity. One possible explanation is that very large apartment buildings have more effective systems for collecting and storing waste away from rats, although other factors such as relatively high socioeconomic status or recent construction of the buildings may play a role. Additionally, these areas may actually harbor many rats but not be good corridors for gene flow, or may be associated with genetic breaks in the rat population for other reasons. Municipal pest management professionals could take advantage of our zoning model to better target scarce resources for rat management to areas where they are most needed. These models also indicate areas where rats are most likely to repopulate through dispersal if high-connectivity areas are not addressed strategically.

The single best predictor of rat genetic connectivity in our modeling was the presence of rail lines, which are overwhelmingly underground subway lines in Manhattan. The first underground subway line opened in Manhattan in 1904 and was an immediate success; ridership nearly doubled in the first decade. Today, the NYC subway has 472 stations and over 1,000 km of track, and is the busiest public transit system in North America by far. A single line running through Manhattan (Lexington Avenue line, including the 4 / 5 / 6 trains) carries more daily passengers (1.3 million daily weekday riders in 2015) than many cities’ entire systems combined. When we sampled rats for this study, daily ridership was historically high at 5.65 million passengers per weekday, but decreased by more than half during the COVID-19 pandemic. Ridership has recovered to 75% of its pre-pandemic ridership as of late 2024^75^. Human consumption of food is a widespread practice on the NYC subway, and thus at a local level subway stations may provide a major infusion of resources to rats. Littering on the platforms and tracks, and subsequent track fires, are a chronic problem in the NYC subway^76^. Many stations also have unsecured trash cans and trash rooms that sustain large local rat populations^10^.

Very little information on pest population dynamics or movements in underground transit systems is available, but our results suggest that they may be important for sustaining rat populations and promoting movement over longer distances than other types of infrastructure. Many subway stations are relatively shallow, and thus rats may move up and down vertically to take advantage of food resources or seek shelter from inclement weather. Rats may also move through subway tunnels under their own power or by riding on trains (personal observation of authors), allowing for rapid unimpeded movements over long distances, which would require traversing more dangerous and complex landscape if attempted aboveground. Some rats that settle in deeper stations may be permanent residents that utilize a particularly valuable resource such as an unsecured trash room. This work advances our understanding of these dynamics using dual population genetic studies: our main landscape genetic modeling, and additional sampling within subway stations and at the surface to explicitly examine genetic relatedness among subway rats.

At very short distances, there was a wide range of genetic distance values between pairs of rats in the subway and at the surface, but particularly at the lowest values. This finding is consistent with there being relatively closely related rats within subway stations, but also with rats at the surface. Thus, there is likely to be at least some movement of rats between subway stations and surface habitats. At distances up to ∼2.8 km, genetic distance is generally lower between pairs of rats in the subway vs pairs of rats at the surface. This finding indicates a greater likelihood of sampling related rats in subway stations, and also suggests that rats are moving between stations and reproducing. Taken as a whole, gene flow with surface populations followed by relatively long-distance movements between subway stations (occasionally facilitated by hitching rides on trains but also moving under their own power through tunnels) indicates that the subway system plays an important role in maintaining long-distance connectivity of the Manhattan rat population. As mentioned previously, typical rat connectivity at the surface is generally limited to a few hundred meters^32^. Subways were also strong predictors in our modeling in combination with zoning and sewer systems, although sewers alone were not a highly ranked variable.

Manhattan, especially in its southern neighborhoods, has extensive sewer lines built of brick that are old and anecdotally known to harbor large rat populations. Old clay or brick sewers near areas of high human population density and food resources are associated with rat infestations^77^, and it’s likely that many of these variables interact with each other and with the presence of underground subway lines.

We also generated a habitat suitability model (HSM) for Manhattan for our landscape genetic analyses. Our HSM is likely the most robust available for *Rattus norvegicus* in an urban context, or perhaps any context, given that the model incorporated thousands of inspection points that were systematically collected by trained municipal officials. Areas with higher human population density, older buildings, and lower median income were associated with active rat signs, which are results that are largely concordant with previous statistical analyses^40,41^. These results broadly make sense given that high human population density defines urban areas^78^, and brown rats are a quintessentially urban commensal. A weakness of our HSM is that there were not systematic surveys conducted in park lands. Anecdotally from trapping for this study, rats are common in edge habitats but generally absent from forested areas in city parks except along water courses. While our HSM outperformed geographic distance and some single variable landscape genetic models, it was still relatively low-ranked among all of the models that we optimized using ResistanceGA. HSMs are now routinely used to parameterize cost or resistance surfaces for landscape genetics and may perform better than expert modeling^79^. However, their performance varies depending on which HSMs are used^80^ and by ecological and behavioral contexts^81^. The species presence data points used to construct HSMs represent a snapshot in time and space for an individual organism, and rats may respond quickly to changing resources or disturbance by dispersing. Landscape genetic modeling in contrast examines how gene flow and genetic drift act over longer periods of time, and part of the signal may accrue over multiple generations as individuals disperse and spread their alleles across an urban landscape. Thus, while HSMs may identify high-quality habitat for rats, they do not necessarily best describe the spatial ecological variation that is associated with rat movements and gene flow. High-quality habitat may actually result in relatively low dispersal rates because resources can be obtained readily at the local scale. Some species readily cross unsuitable habitat during dispersal events and thus HSMs would not adequately capture these dispersal events^81^. Urban rats may move several km when released in unfamiliar territory^82^. While rare, such events may influence their population genetic structure. Species’ movements may also vary over time due to the influence of humans or other disturbances^83^. The optimization approach we used here to fit pairwise genetic distances to landscape resistance values^51^, rather than relying on *a priori* resistance assignments, has also previously been shown to outperform HSMs^84^. While resistance-based landscape genetic modeling has proven very useful here and in many other studies, future advances will incorporate more realistic scenarios of spatiotemporal variation in animal movement^83^.

In this paper we bring together landscape connectivity, population genetic structure, and urbanization in a unique analysis of an iconic, economically important urban commensal species. The brown rat has a rich history of association with humans^37^, and rats will be an important component of urban areas moving forward as the human population becomes increasingly concentrated in cities. New approaches to incorporating ecological, evolutionary, and human social variation^26^ into analyses of non-human species hold much promise for promoting desired species and limiting harmful species in cities. Our findings that subways, zoning regulations, and social factors influence habitat suitability and genetic connectivity of an urban rat population indicate that management of urban biodiversity requires multifaceted approaches.

## Data availability

DNA sequencing reads for this project are available on the NCBI SRA at accession PRJNA414893. Resistance layers and habitat suitability maps will be shared upon request.

## Code availability

The R code for the ResistanceGA modeling conducted in this study is available upon request and will be made available on Github.

## Acknowledgements

We thank Sonia Bernal, David Forlano, Brittney Kajdacsi, Nicholas Rapillo, Matt Wollman and Otis Wood, and the Ryders Alley Trencher-Fed Society (R.A.T.S.) for help with sample collection, and Jane Park and Ian Hayes for laboratory work support. Jonathan Richardson provided many thoughtful comments early in this project. We thank the City of New York Department of Parks and Recreation for trapping permits, the Department of Health and Mental Hygiene for access to rat-related datasets, and the Metropolitan Transportation Authority for assistance with field work in the subway system.

## Author contributions

Conceptualization: J.M.-S., M.C.; Investigation: J.M.-S., M.C.; Writing – original draft: J.M.-S., M.C.; Writing – review and editing: J.M.-S., M.C.; Visualization: M.C.; Supervision: J.M.-S.; Project administration: J.M.-S.

## Supplementary information The PDF file includes

Supplemental Methods Figs. S1-S8

Tables S1-S2

