## Supplementary Materials for "Subways, zoning and social factors influence the dispersal and genetic connectivity of an iconic urban pest, the brown rat (*Rattus norvegicus*) across a complex cityscape"

Supplemental Methods: Landscape variable generation and data sources.

### 1. Non-Built Physical Factors

*Topographical Slope*: We predicted that high values of topographical slope might limit rat habitat suitability and gene flow, while lower values (gradual changes in elevation) might have an opposite effect by providing burrowing opportunities[1] .We used available digital elevation maps to calculate percent slope at the finest possible resolution (i.e. 1m), then resampled to the final 30m resolution using bilinear interpolation in ARCMAP.
Topographical Slope data source: https://www.usgs.gov/core-science-systems/ngp/tnm-delivery/

*Green Landcover*: We expected that green space would increase habitat suitability and promote rat gene flow within urban landscapes[2,3]. Using available landcover datasets we isolated green cover into binary categorical datasets for use in landscape genetic analysis.
Green Landcover data source: https://opendata.cityofnewyork.us/

*Percentage Green Space*: Because rats use habitat and resources within their home range and across dispersal corridors, we included a variable representing the percentage of green space within a defined radius. Using an available 1m$^2$ landcover map, we calculated the percentage of green land cover within a 100m radius of each raster cell for habitat suitability modeling and within a 400m radius of each raster cell for landscape genetic analysis, then resampled to 30m$^2$ using the bilinear method. The difference in radius used was due to different expectations of use as home range habitat (which is spatially limited in cities) and dispersal (which likely extends far past the normal home range).
Percentage Green Space data source: https://opendata.cityofnewyork.us/

*Dirt Landcover*: We expected that exposed soil would increase habitat suitability and promote rat gene flow within urban landscapes[2,3]. Using available landcover datasets we isolated dirt landcover into binary categorical datasets for use in landscape genetic analysis.
Dirt Landcover data source: https://opendata.cityofnewyork.us/

*Water Landcover*: Rats require daily access to fresh water for drinking. Waterways have been shown to restrict movement[4], though banks of waterways also provide valuable habitat and movement corridors for rats[1]. Using available landcover datasets we isolated water landcover into binary categorical datasets for use in landscape genetic analysis.
Water Landcover data source: https://opendata.cityofnewyork.us/

### 2. Built Physical Factors

*Impervious Landcover*: Paved surfaces can prevent burrowing by rats and traffic on roads can be a source of mortality and disturbance that might impede rat movement. Using available landcover datasets we isolated impervious landcover not classified as buildings into binary categorical datasets for use in landscape genetic analysis.
Impervious Landcover data source: https://opendata.cityofnewyork.us/

*Built Landcover*: Built structures can provide sources of harborage and resources for rats due to structural gaps, lack of maintenance, and anthropogenic resources. Using available landcover datasets we isolated landcover classified as buildings into binary categorical datasets for use in landscape genetic analysis.
Built Landcover data source: https://opendata.cityofnewyork.us/

*Subway Tunnels*: Subway tunnels provide long stretches of simple underground habitat that link aboveground locations. Rail lines are known to provide movement corridors for urban wildlife[5] Rats are known to inhabit subway stations, which provide anthropogenic resources and climate stability. For habitat suitability modeling, we created rasters of Euclidean distance to rail lines based on available polyline shapefiles. For landscape genetic modeling we made categorical rasters of rail line presence.
Subway Tunnel data source: https://opendata.cityofnewyork.us/

*Building Age*: Older built structures have been found to be more conducive to rat habitat[6,7], which may also impact capcacity for rat movement. Older structures may be less structurally sound, experience more renovations impacting utilities and other building penetrations, and exhibit lower overall upkeep.We used the PLUTO database, which provides building specific information on each tax lot including the age of the building. We built a raster of building age at 1m, then filled pixels representing parks with the median value (1900) and filled pixels representing streets with the average age of nearby buildings using the focal statistics tool from the Spatial Analyst Toolbox in ARCMAP. The resulting raster was used for habitat suitability and landscape genetic modeling.
Building Age data source: https://www.nyc.gov/site/planning/data-maps/open-data/dwn-pluto-mappluto.page

*Roadway Width*: Increased road traffic on larger roads may limit rat movement through mechanisms of increased mortality and behavioral avoidance[8,9]. Roadway width is a useful proxy for road traffic in NYC. We used spatial data on roadway width to generate a categorical layer of minor roads (0 - 52ft wide), major roads (52 - 85ft wide), or non-road areas based on natural breaks in the overall distribution of road widths. Major roads captured most avenues (North-South direction) and larger two-way streets (East-West direction), while minor roads captured most other one- way streets in Manhattan.
Roadway Width data source: DCM_StreetCenterLine from https://www.nyc.gov/site/planning/data-maps/open-data/dwn-digital-city-map.page

*Brick Sewer Density*: Brown rats are known to occupy sewer infrastructure in cities and move greater distances through sewer networks than through above ground urban matrix on average[10,11]. We expected rats to preferentially use older sewers and sewers that are composed of brick building materials, as compared to newer sewers built with poured cement[12]. In NYC, we used spatial data describing the building material of catch basins (street entrances to sewers) to isolate those made of brick. We calculated the kernel density of brick sewers using a 100m search radius for use in habitat suitability modeling, and with a 400m search radius for landscape genetic analysis. The difference was due to different expectations of use as home range habitat

(which is spatially limited in cities) and dispersal (which likely extends far past the normal home range).
Brick Sewer data source: NYC Department of Health and Mental Hygiene (data available from authors upon request)

*Restaurant Density*: Urban rats exploit anthropogenic food resources. We predicted that areas where humans have developed higher densities of food service establishments (i.e., restaurants and food carts), we would observe increases in rat habitat suitability and gene flow. Using available point location data on the presence of food service establishments, we generated a kernel density layer using a 400m radius around each 1m raster cell, then resampled to the final 30m resolution using the bilinear method.
Restaurant Density data source: NYC Department of Health and Mental Hygiene (data available from authors upon request)

3. Social Factors

*Population Density*: Brown rats are highly associated with human presence, so we expected rat habitat suitability and gene flow to increase with increasing density of humans[13]. Using public census data at the census block level, we divided the number of residents within each block by the area (square kilometers) to calculate population density.
Population Density data source: https://www.census.gov/geo/maps-data/data/tiger-data.html

*Median Income:* Lower income areas in cities have been shown to support higher quality rat habitat[13,14]. We used census data at the census block level to generate a raster of median annual income using data from the American Community Survey, which averages data over a five-year period. We used a data spanning 2011-2016, which overlaps our rat sampling period.
Median Income data source: https://www.census.gov/geo/maps-data/data/tiger-data.html

*Business Improvement Districts*: NYC has created business improvement districts (BIDs), where business owners collectively support efforts to improve sanitation (e.g., through daily street litter collection). These areas may exhibit reduced rat habitat and gene flow through a reduction in resources and increase in overall maintenance. Using available shapefiles of BID boundaries, we generated categorical rasters of BIDs.
BIDs data source: https://opendata.cityofnewyork.us/

*Municipal Zoning*: We predicted that geographical areas with different zoning designations would have differential effects on rat habitat suitability in gene flow. We hypothesized that higher density residential areas supporting greater habitat and movement, while lower density and industrial/manufacturing areas would support lower quality habitat and less movement. Using available spatial data, we grouped zoning designations into several categories of similar types to simplify interpretation, resulting in 9 categories (1: Low/Medium density residential, 2: High density residential, 3: Very high density residential, 4: Commercial for local use, 5: Commercial for regional use, 6: High bulk commercial areas, 7: Light manufacturing, 8: Medium/Heavy manufacturing, 9: Park or playground).

NYC data source: https://data.cityofnewyork.us/City-Government/Zoning-GIS-Data-Shapefile/kdig-pewd

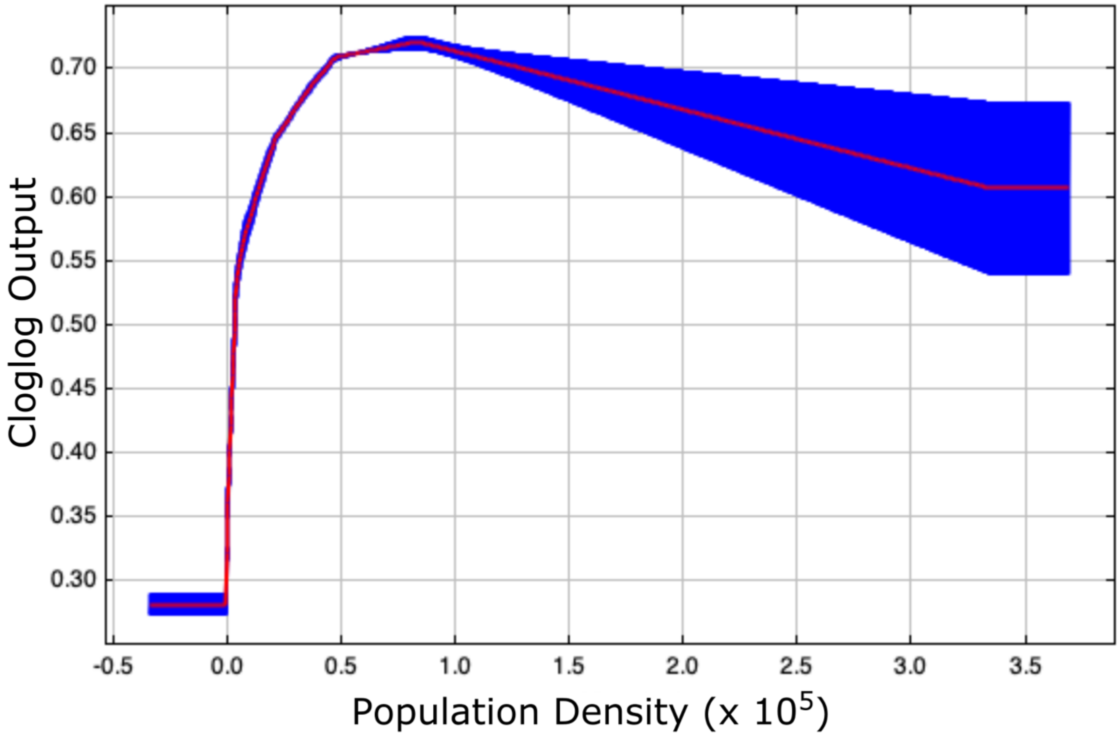


Supplementary Figure 1: Maxent response curve for population density, which provided 33.1% contribution towards the final habitat suitability model.

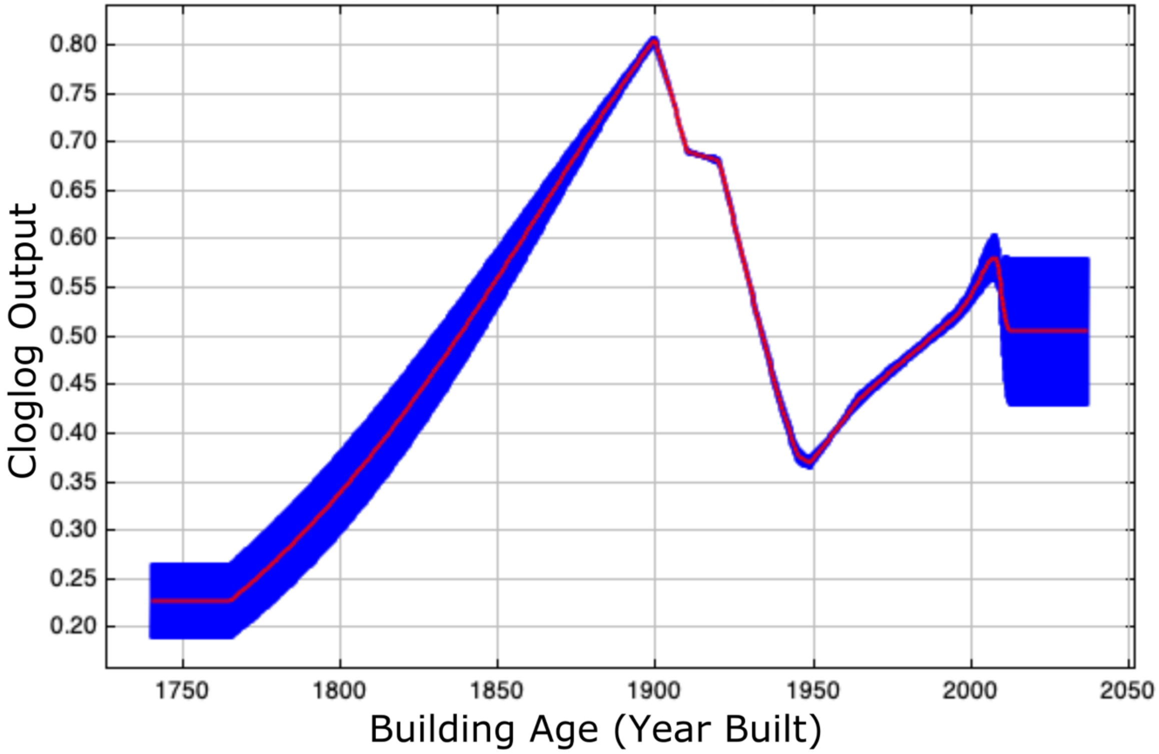


Supplementary Figure 2: Maxent response curve for building age, which provided 20.4% contribution towards the final habitat suitability model.

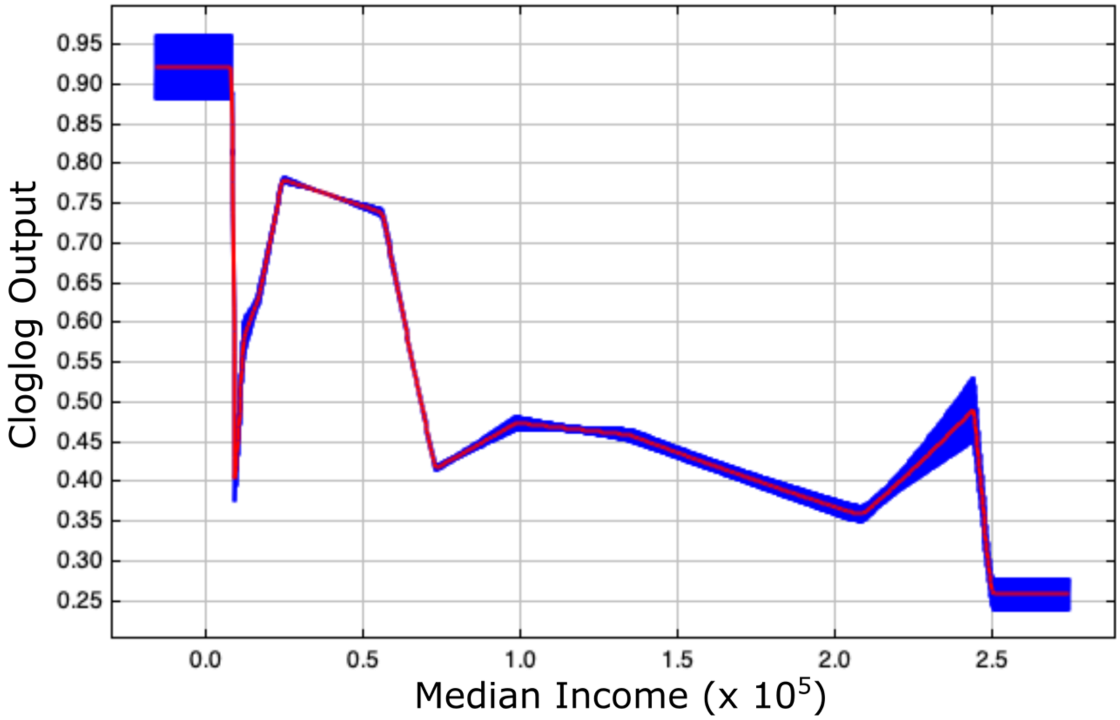


Supplementary Figure 3: Maxent response curve for median income, which provided 16.0% contribution towards the final habitat suitability model.

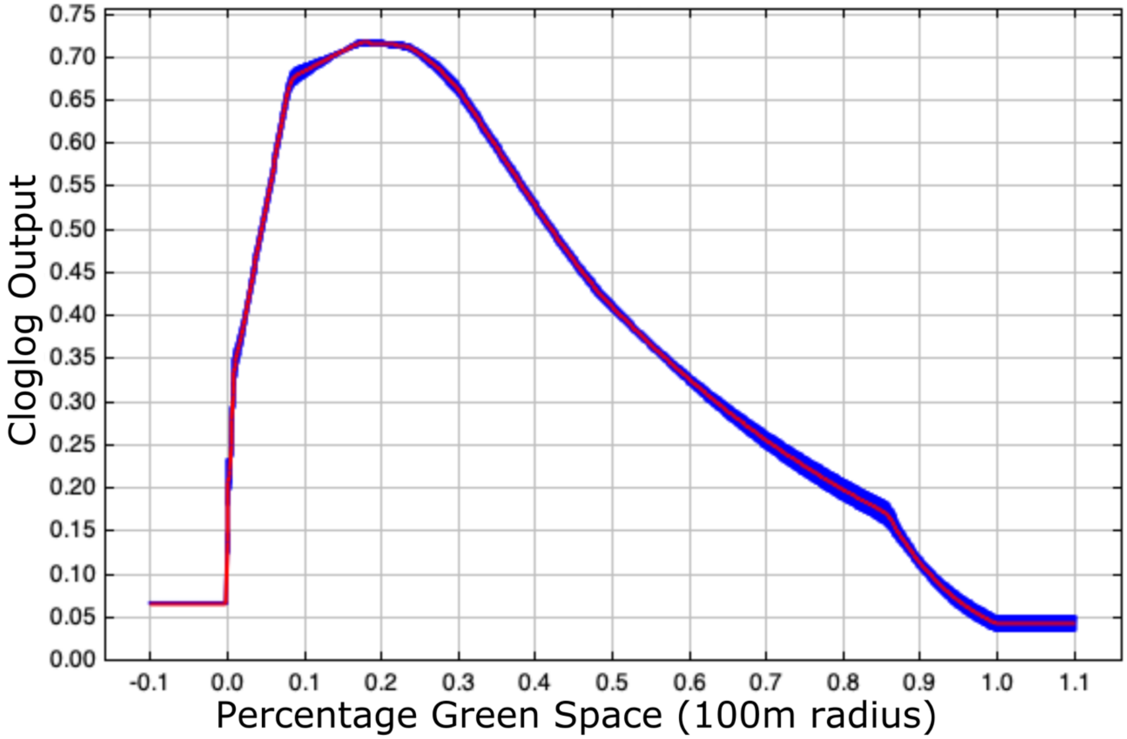


Supplementary Figure 4: Maxent response curve for the percentage of local green space within a 100m radius, which provided 9.9% contribution towards the final habitat suitability model.

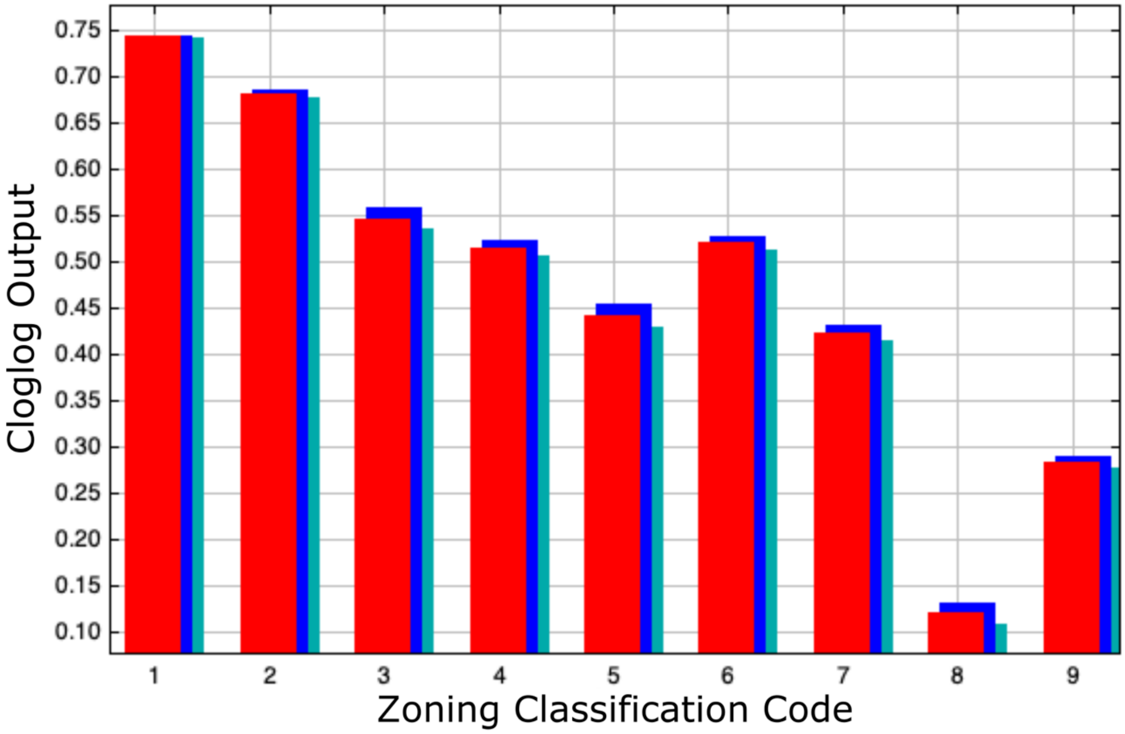

Supplementary Figure 5: Maxent response curve for zoning classification, which provided 8.1% contribution towards the final habitat suitability model. 1: Low/Medium density residential, 2: High density residential, 3: Very high density residential, 4: Commercial for local use, 5: Commercial for regional use, 6: High bulk commercial areas, 7: Light manufacturing, 8: Medium/Heavy manufacturing, 9: Park or playground.

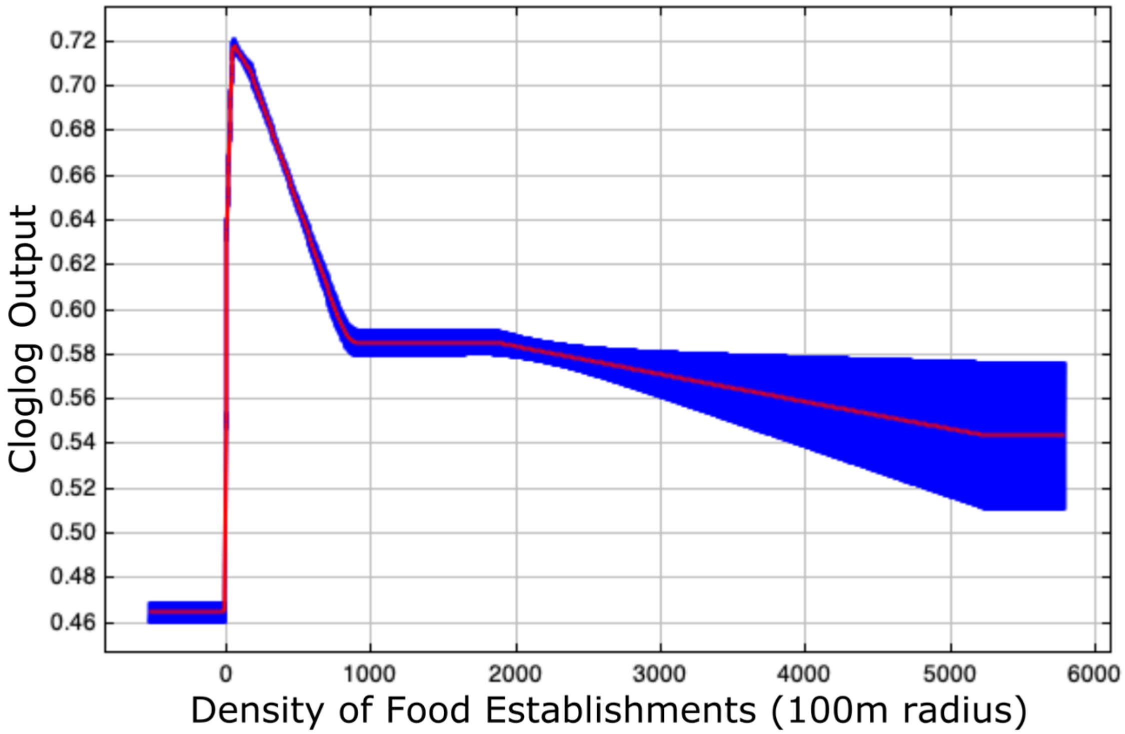


Supplementary Figure 6: Maxent response curve for the kernel density of human food establishments within a 100m radius, which provided 4.8% contribution towards the final habitat suitability model.

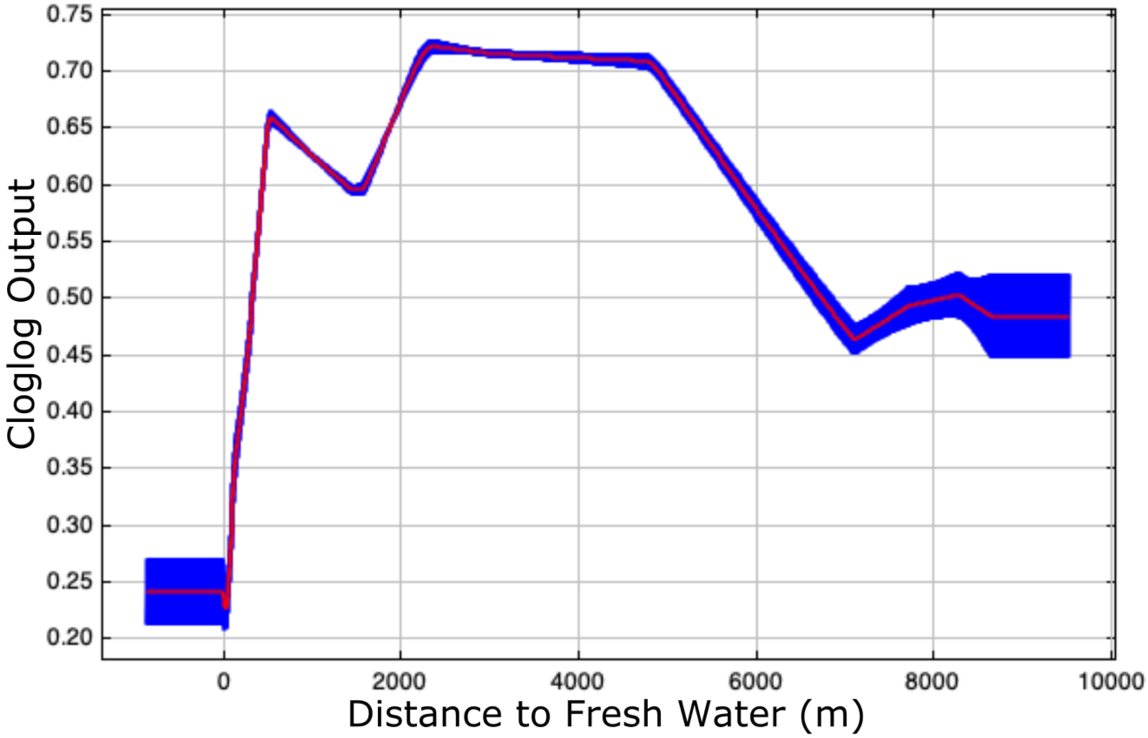


Supplementary Figure 7: Maxent response curve for the distance to fresh water, which provided 4.0% contribution towards the final habitat suitability model.

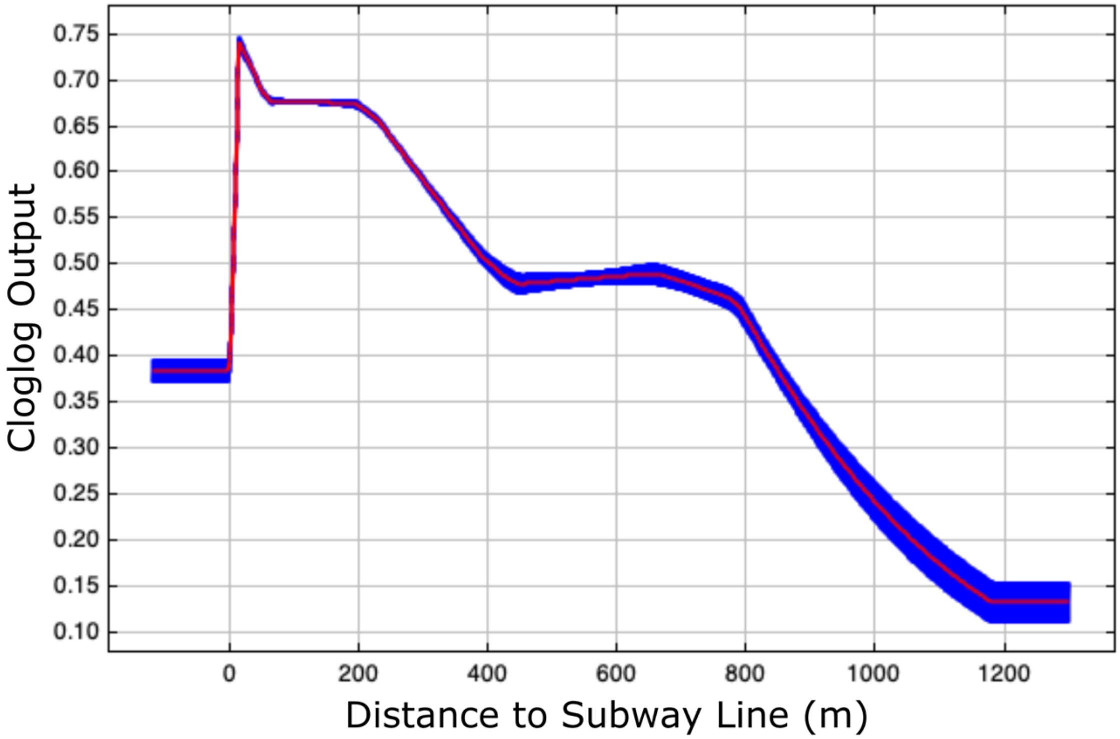


Supplementary Figure 8: Maxent response curve for the distance to a subway line, which provided 3.8% contribution towards the final habitat suitability model.

| Multivariate Model | Variables Included | Rationale |
|---|---|---|
| Subway & Sewer | Subway Tunnels, Sewer Material | Combined effect of underground infrastructure |
| Subway & Zoning | Subway Tunnels, Municipal Zoning | Combined effect of movement through subways and variable zoning classifications |
| All Social | BIDs, Median Income,Municipal Zoning, Population Density, Restaurant Density | Combined effect of all social factors |
| Rat Food | Green Landcover, Population Density,Restaurant Density | Combined effect of areas providing obvious food resources |
| All Natural | Dirt Landcover, Green Landcover, Topigraphical Slope, Water Landcover | Combined effect of all nonbuilt physical factors |
| GreenLC/PopDen | Green Landcover, Population Density | Combined effect of habitat and food resources in green areas and impacts of variable population density |
| Landcover | Built Landcover, Dirt Landcover, Green Landcover, Impervious Landcover, Water Landcover | Combined effect of full landcover matrix |
| Natural Resources | Green Landcover, Water Landcover | Comined effect of naturally occuring resources |
| GreenLC/Income | Green Landcover, Median Income | Combined effect of habitat and food resources in green areas and impacts of variable neighborhood income |
| PercentGreen/PopDens | Percentage Green Space, Population Density | Combined effect of neighborhood scale green space availability and variable population density |
| All Built | Building Age, Built Landcover, Impervious Landcover, Street Width, Sewer Material, Subway Tunnels | Combined effect of all built physical factors |
| PercentGreen/Income | Median Income, Percentage Green Space | Combined effect of neighborhood scale green space availability and variable neighborhood income |
| Income/PopDens | Median Income, Population Density | Combined effect of variable neighborhood income and populatio density |

| Iteration | Type | Model | obj.func_LL | k | AIC | AICc | R2m | R2c | LL | AICcRank |
|---|---|---|---|---|---|---|---|---|---|---|
| 2 | Uni | Subway Tunnels | 79910.4731 | 3 | -159814.95 | -159814.85 | 0.59361956 | 0.96384044 | 79910.4731 | 1 |
| 2 | Multi | Subway & Sewer | 79810.421 | 6 | -159608.84 | -159608.51 | 0.58369386 | 0.95916134 | 79810.421 | 2 |
| 2 | Uni | Municipal Zoning | 78261.4544 | 10 | -156502.91 | -156502.03 | 0.64622427 | 0.9719923 | 78261.4544 | 3 |
| 2 | Multi | Subway & Zoning | 78246.9726 | 12 | -156469.95 | -156468.69 | 0.53989853 | 0.9307979 | 78246.9726 | 4 |
| 2 | Multi | All Social | 76047.9129 | 21 | -152053.83 | -152049.98 | 0.47868401 | 0.87857346 | 76047.9129 | 5 |
| 2 | Multi | Rat Food | 75493.4405 | 9 | -150968.88 | -150968.17 | 0.61935946 | 0.96849389 | 75493.4405 | 6 |
| 2 | Multi | GreenLC & PopDen | 75464.1097 | 6 | -150916.22 | -150915.89 | 0.61420647 | 0.97330877 | 75464.1097 | 7 |
| 2 | Multi | All Natural | 75390.2297 | 10 | -150760.46 | -150759.58 | 0.57669846 | 0.95277593 | 75390.2297 | 8 |
| 2 | Uni | Roadway Width | 75217.1967 | 4 | -150426.39 | -150426.24 | 0.58340869 | 0.95467898 | 75217.1967 | 9 |
| 2 | Multi | Landcover | 75193.9856 | 11 | -150365.97 | -150364.92 | 0.60942337 | 0.96948937 | 75193.9856 | 10 |
| 2 | Uni | Distance to Earthen Space | 75092.2617 | 4 | -150176.52 | -150176.37 | 0.51083249 | 0.91983852 | 75092.2617 | 11 |
| 2 | Multi | Natural Resources | 75048.4905 | 5 | -150086.98 | -150086.75 | 0.61572612 | 0.97105411 | 75048.4905 | 12 |
| 2 | Uni | Green LC | 75046.2132 | 3 | -150086.43 | -150086.33 | 0.61572844 | 0.97098596 | 75046.2132 | 13 |
| 2 | Multi | GreenLC & Income | 75046.2132 | 6 | -150080.43 | -150080.1 | 0.6157289 | 0.97098621 | 75046.2132 | 14 |
| 2 | Uni | Percentage Green Space | 74859.6849 | 4 | -149711.37 | -149711.21 | 0.45525327 | 0.89465971 | 74859.6849 | 15 |
| 2 | Multi | All Built | 74827.034 | 16 | -149622.07 | -149619.85 | 0.46101283 | 0.8712896 | 74827.034 | 16 |
| 2 | Multi | PercentGreen & PopDens | 74568.0716 | 7 | -149122.14 | -149121.7 | 0.48495721 | 0.90932106 | 74568.0716 | 17 |
| 2 | Multi | PercentGreen & Income | 74550.0529 | 7 | -149086.11 | -149085.66 | 0.42086083 | 0.87626651 | 74550.0529 | 18 |
| 2 | Uni | Sewer Material | 74286.8255 | 4 | -148565.65 | -148565.5 | 0.52144105 | 0.89530602 | 74286.8255 | 19 |
| 2 | Uni | Built LC | 74135.0619 | 3 | -148264.12 | -148264.03 | 0.52628708 | 0.91032729 | 74135.0619 | 20 |
| 2 | Uni | Topographic Slope | 73884.9395 | 4 | -147761.88 | -147761.72 | 0.26596818 | 0.77964784 | 73884.9395 | 21 |
| 2 | Uni | Impervious Surface LC | 73877.4604 | 3 | -147748.92 | -147748.83 | 0.52137958 | 0.93045566 | 73877.4604 | 22 |
| 2 | Uni | Building Age | 73794.4934 | 4 | -147580.99 | -147580.83 | 0.61176136 | 0.96864641 | 73794.4934 | 23 |
| 2 | Uni | Population Density | 73768.5031 | 4 | -147529.01 | -147528.85 | 0.37560597 | 0.82285716 | 73768.5031 | 24 |
| 2 | Uni | Habitat Suitability Model | 73613.3412 | 4 | -147218.68 | -147218.53 | 0.53637937 | 0.92338098 | 73613.3412 | 25 |
| 2 | Multi | Income & PopDens | 73591.1645 | 7 | -147168.33 | -147167.89 | 0.23261686 | 0.74042962 | 73591.1645 | 26 |
| 2 | Uni | Restaurant Density | 73358.7642 | 4 | -146709.53 | -146709.37 | 0.48993889 | 0.86355908 | 73358.7642 | 27 |
| 2 | Uni | BIDs | 73260.9658 | 3 | -146515.93 | -146515.84 | 0.54430398 | 0.90948685 | 73260.9658 | 28 |
| 2 | Uni | Water LC | 72843.9135 | 3 | -145681.83 | -145681.73 | 0.16500779 | 0.70902729 | 72843.9135 | 29 |
| 2 | Uni | Median Income | 72828.2558 | 4 | -145648.51 | -145648.36 | 0.17288169 | 0.71450126 | 72828.2558 | 30 |
| 2 | Uni | Dirt LC | 72822.2787 | 3 | -145638.56 | -145638.46 | 0.19959094 | 0.73763149 | 72822.2787 | 31 |
| 2 | Uni | Distance | 72789.8202 | 2 | -145575.64 | -145575.59 | 0.16056309 | 0.70617504 | 72789.8202 | 32 |
